# Activation of Shh/Smo is sufficient to maintain oligodendrocyte precursor cells in an undifferentiated state but is not necessary for differentiation

**DOI:** 10.1101/2023.06.23.546285

**Authors:** Sonia Nocera, Miguel A. Marchena, Beatriz Fernández-Gómez, Yolanda Laó, Christian Cordano, Óscar Gómez-Torres, Rafael Luján, Fernando de Castro

## Abstract

Myelination is the terminal step in a complex and precisely timed program that orchestrates the proliferation, migration and differentiation of oligodendroglial cells. It is thought that Sonic Hedgehog (Shh) acting on Smoothened (Smo) participates in regulating this process, but that these effects are highly context dependent. Here, we investigate oligodendroglial development and remyelination from three specific transgenic lines: NG2-Cre^ERT2^ (control), Smo^fl/fl^/NG2-Cre^ERT2^ (loss of function) and SmoM2/NG2-Cre^ERT2^ (gain of function), as well as pharmacological manipulation that enhance or inhibit the Smo pathway (SAG or cyclopamine treatment respectively). To explore the effects of Shh/Smo on differentiation and myelination in vivo, we developed a highly quantifiable model by transplanting OPCs in the retina. We find that myelination is greatly enhanced upon cyclopamine treatment and hypothesize that Shh/Smo could promote OPC proliferation to subsequently inhibit differentiation. Consistent with this hypothesis, we find that the genetic activation of Smo significantly increased numbers of OPCs and decreased oligodendrocyte differentiation when we examined the corpus callosum during development and after cuprizone demyelination and remyelination. However, upon loss of function with the conditional ablation of Smo, myelination in the same scenarios are unchanged. Taken together, our present findings suggest that the Shh pathway is sufficient to maintain OPCs in an undifferentiated state, but is not necessary for myelination and remyelination.

## INTRODUCTION

Oligodendrocytes (OLs) are the myelinating cells in the central nervous system (CNS) and have been identified and studied for over a century (del Río-Hortega, 1919). OLs arise from oligodendrocyte precursor cells (OPCs) during pre- and postnatal development, and the persistent presence of OPCs in the adult CNS and in demyelinating lesions has been extensively investigated (Small *et al*., 1987; Wolswijk & Noble, 1989; Warf *et al*., 1991; Levine *et al*., 2001). Despite this, numerous aspects of the proliferation and differentiation processes of these OPCs during development, demyelination and subsequent remyelination remain unknown (Franklin et al., 2012). Shh is a widely studied factor in relation to the oligodendrocyte lineage. Most of these studies focus on the specification of neural progenitor cells to oligodendrocyte precursor cells (Fuccillo *et al*., 2006; Wang *et al*., 2008; Ferent *et al*., 2013; Mierzwa *et al*., 2014; Samanta *et al*. 2015; Del Giovane & Ragnini-Wilson, 2018; Sanchez & Armstrong, 2018; Winkler *et al*., 2018; Namchaiw *et al*., 2019; Zakaria *et al*., 2019; Macchi *et al*., 2020; Ming *et al*., 2020; Del Giovane *et al*., 2021; Klein *et al*., 2021; Laouarem et al., 2021; Nguyen *et al*., 2021; Del Giovane et al., 2022). Smoothened (Smo) is the main Shh receptor (Petrova & Joyner, 2014; Ruatt et al., 2014), and is especially relevant in the processes mentioned previously but abundant data available to date do not provide mechanistic insight on the role of Shh and Smo in oligodendrogial maturation, instead displaying several contradictions. While Shh is required for the specification of OPCs in ventral oligodendrogliogenic niches during embryonic development (Fuccillo *et al*., 2006), in the adult brain Shh promotes OLs generation (Loulier et al., 2006) and ameliorates EAE clinical course, preventing demyelination (Xiao et al., 2021). In addition, the target genes of the Shh cascade show increased activity during remyelination of adult brain lesions, suggesting a reactivation of the pathway (Wang *et al*., 2008; Ferent *et al*., 2013; Sánchez y Armstrong, 2018). However, other studies show that inhibition of the Shh pathway results in OLs maturation and remyelination (Samanta *et al*., 2015; Namchaiw *et al*., 2019; Radecki *et al*., 2020). To add even more complexity, several studies suggest that the Shh pathway can interact with other elements of the molecular environment during development or during pathological conditions (Kessaris *et al*., 2004; de Castro & Bribián, 2005; Furusho *et al*., 2011; de Castro & Zalc, 2020; Macchi et al., 2020; Ming et al., 2020).

Our present work explores the role of Shh and the Smo receptor in different contexts: i) early postnatal development, ii) oligodendroglial dynamics during active demyelination, and iii) (re)myelination. It is easy to think that these three conditions possess very different cellular and molecular “microenvironments” from each other. Moreover, it is likely that these microenvironments underlie the differences found between the contrasting reports about Shh functions. Therefore, prior to investigating these context-dependent roles for Shh, we developed an *in vivo* method for testing compounds for their ability to enhance OPC differentiation and myelination using retinal ganglion axons in the retina--which are normally unmyelinated due to limited access to oligodendroglial cells (Setzu *et al*., 2006; Yuen *et al*., 2013). We decided to use it without combining Zimosan-induced inflammation to study the physiologically non-myelinated intraretinal axonal segments of the retinal ganglion axons in an environment completely free of any developmental or reparative influence (chemotropic molecules, growth factors, cytokines, etc.) which make it an ideal platform to study myelination and the pathways that influence this complex process. In this highly relevant in vivo model, we find that myelination is greatly enhanced upon cyclopamine treatment and hypothesize that Shh/Smo could promote OPC proliferation to subsequently inhibit differentiation. Consistent with this hypothesis, the constitutive activation of Smo with the SmoM2/NG2-Cre^ERT2^ mice significantly increased numbers of OPCs and decreased oligodendrocyte differentiation during development of the corpus callosum and after cuprizone demyelination and remyelination. However, upon loss of function with the conditional ablation of Smo (Smo^fl/fl^/NG2-Cre^ERT2^), myelination is unaffected. Taken together, our present findings suggest that Shh pathway is sufficient to maintain OPCs in an undifferentiated state but is not necessary for myelination and remyelination.

## METHODS

### Animals

All animal experiments were approved by the local ethics committee CSIC (440/2016, Madrid, Spain) and the Spanish and European regulations (RD 53/2013 and 178/2004, Ley 32/2007 y 9/2003, Decree 320/20102010/63/EU, 90/219/EEC). Mice were housed at the Animal facilities of the Instituto Cajal on a 12 h light/dark cycle with ad libitum access to food and water. NG2-Cre tdTomato mice were gently donated by Kirchhoff lab (University of Saarland, Homburg, Germany). The targeting strategy used for his creation is described by Huang et al., 2014. Conditionally Smo floxed and R26SmoM2 mice were purchased at JAX lab (#004526, Long et al. 2001; #005130, Jeong et al. 2004). R26SmoM2 line contains a copy of the Smoothened gene inserted into the Gt(ROSA)26Sor locus. The mutant allele contains a point mutation W539L, which renders it constitutively active. The conditional NG2-Cre^ERT2^;Rosa26tdTomato line was crossbred with SmoM2 and Smo floxed lines. Recombination was induced with tamoxifen at P1-P2, and the effect of Shh modulation was examined at P14 when OPCs and OLs populate the CC during postnatal myelination (Kessaris et al. 2006; Sanchez e Armstrong 2018).

### Tissue preparation

Mice were transcardially perfused with 4% PFA, brains postfixed in 4% PFA during 4h at RT, washed in PB (0.1 M), and cryoprotected through increasing concentration (w/v) of sucrose. For immunohistochemistry, brains were sectioned coronally at 30 (P14) and 20μm (adult) with sliding microtome and cryostat, respectively.

### Immunostaining

Samples were pre-incubated with blocking solution (5% normal donkey serum and 0.2% Triton X-100 in PBS). Incubation with primary and secondary antibodies was carried out overnight at 4°C and for 2h at RT, respectively. Primary antibodies used were anti-PDGFRα 1:150 (goat, R&D system), anti-CC1 1:100 (mouse, Millipore), anti-MBP 1:500 (Rat, Bio-Rad), anti-Olig2 1:200 (Rabbit, Millipore), anti-CASPR 1/900 (Rabbit, Abcam); and Alexa Fluor-conjugates secondary antibodies, 1:1000 (Invitrogen).

### Tamoxifen administration

Tamoxifen (Sigma-Aldrich) was dissolved in corn oil. Transgenic mice received tamoxifen at P1-2 by the lactating mothers (one intraperitoneal injection of 100 mg/kg body weight), and the brains were perfused at P14. Tamoxifen (100 mg/kg body weight) was administered intraperitoneally once per day for five consecutive days for young adult mice. No labeling was seen in absence of tamoxifen administration.

### Cuprizone demyelination model

As previously in our group (Medina-Rodríguez et al., 2017), 8-9-week-old mice fed a diet of chow mixed with 0.25% cuprizone (Sigma-Aldrich) ad libitum integrated into a powdered standard rodent chow. Toxic demyelination was induced with a five weeks Cuprizone diet (42 days in total) in 4 groups of mice (at least three males for group). After five weeks, mice were back to the standard chow. Tamoxifen was given in different strategical periods for five consecutive days. The first group received it at the beginning of the cuprizone diet. Mice were perfused after week 3 to test OPCs proliferation and microglia activation. Tamoxifen was also given at week 5, and mice perfused after two weeks of recovery to test oligodendrocyte differentiation and remyelination. Aged-matched control mice with a regular diet were taken from each group for immunohistochemical analysis.

### EdU detection

5-ethynyl-2’-deoxyuridine (EdU, Invitrogen) was dissolved at 0.2 mg/ml in the drinking water to detect proliferating cells. The water was exchanged every 48h to 72h, and mice were perfused after seven days of EdU exposure. For the EdU visualization, sections were rinsed with 100mM Tris HCl pH 8.5 immediately after immunohistochemistry and incubated with a reaction solution composed of 100mM Tris HCl pH 8.5 (10%), 100mM CuO4 (2%), Alexa488 azide 2nM (0.5%), dissolved in mqH2O. Finally, ascorbic acid (10%) was added before adding the solution to the sample. After 30 minutes at RT, sections were washed and mounted with a coverslip in Fluoromount-G.

### OPC Culture and intraretinal injection

Tamoxifen induction was made to lactating mothers at P2, and P7 transgenics mice were used for each primary culture. OPCs purification was made through magnetic-activated cell sorting, MACS using O4 antibody (Dincman et al. 2012; Bribián et al., 2020). Adult mice were anesthetized with xylazine/ketamine (87.5mg/kg Ketamine/12.5 mg/kg Xylazine) through intraperitoneal injection (IP). Cells were injected intravitreally, under direct visual observation, close to the RGC layer with a 33-gauge needle and 5μL glass syringe (Hamilton). OPCs were delivered together with the 1μL of cell suspension. Animals were perfused after five weeks. Injected eyes were removed and placed in 4% PFA for 2 hours, then transferred to PBS. Retinae were dissected from the sclera for immunohistochemistry. Finally, after making 4-5 radial incisions, flat-mounted retinae were mounted on slides and examined under confocal microscopy.

### Confocal microscopy

Images were acquired with an inverted Leica SP5 confocal microscopy. Microphotographs were captured at 20X magnification with a z-plane depth of 3 μm or 40X with a z-plane depth of 1 μm. For CASPR-MBP double-staining, a 63X objective was used, microphotographs were taken in a single z-stack with the major MBP intensity. Confocal images were processed using the Image J processing package FIJI (Schindelin et al. 2012). Cell count was performed manually. The rostral CC was analyzed between the Bregma 1.10 and 0.14 μm. Regions of interest (ROIs) were defined from the midline to the point under the peak of the cingulum for P14 experiments or the center of the CC for the cuprizone model.

### Electron microscopy

Animals were transcardially perfused first with 0,9% NaCl, followed by a fixative solution containing 1% glutaraldehyde and 4% PFA in 0.1 M PB (pH 7.4). Brains were postfixed overnight at 4°C, then washed with PB for two hours. 60 μm-thick coronal sections were obtained using a vibrating microtome. Tissue was postfixed with 1% OsO4 in PB for 30 minutes, counterstained with 1% uranyl acetate in dH2O for 30 minutes, and dehydrated with a series of graded ethanol until 100% infiltrated with propylene oxide and embedded in epoxy resin Durcupan. Ultrathin sections of the CC (70 nm-thick sections) were cut using an ultramicrotome, collected in single slot copper grids at RT, and analyzed using a transmission electron microscope at 1500X magnification, as previously reported (Murcia-Belmonte et al., 2016). ImageJ/Fiji was used to obtain axon diameter measurements (Feret diameter, https://imagej.nih.gov/ij/docs/menus/analyze.html). For G-ratio, the inner myelin sheath diameter was divided by the outer myelin sheath diameter. Analyses took at least 500 axons/animal, 3 animals/genotype in the cuprizone group, and 1 animal/genotype in the no-treated group.

### Statistical analysis

Statistics were performed with the GraphPad Prism version 7. Data are expressed as the mean ± the standard error of the mean (SEM). Student’s *t*-test was used to compare two populations/groups and One-Way-ANOVA to compare more than two groups. The threshold for statistical significance was established at p < 0.05. The results of the different statistical analysis are represented in figures as follows: * p < 0.05; ** p < 0.01; and *** p < 0.001.

## RESULTS

### The transplantation of OPCs into the retina is a viable and effective method for studying the role of the Shh-Smo pathway, as well as other molecules, receptors and pathways that influence myelination

To assess the effect of the Shh/Smo pathway in a more simplistic and reduced microenvironment, we exploited the unmyelinated axons of the retina. First, we injected OPCs expressing tdT into the vitreous humor, near the optic nerve head (figure 1A). We verified that the injected OPCs survive in this space (figure 1B). In addition to survival, these cells are able to differentiate and myelinate the axons present in the fiber layer of the optic nerve (figure 1C-D). Having established the feasibility of the injection system and the viability of OPCs in the vitreous humor, we wanted to test the effect of activation or inhibition of the Smo receptor on myelination. To do this, we co-injected OPCs with either cyclopamine (inhibitor) or SAG (activator). Five weeks after transplantation of the OPCs, we were able to observe that the presence of cyclopamine leads to a greater differentiation of the OPCs and, with it, a greater myelinated area (figure 1E-G, K). However, the presence of SAG (activator) did not alter the myelinated area with respect to the controls (figure 1H-J, K).

**Figure 1.**
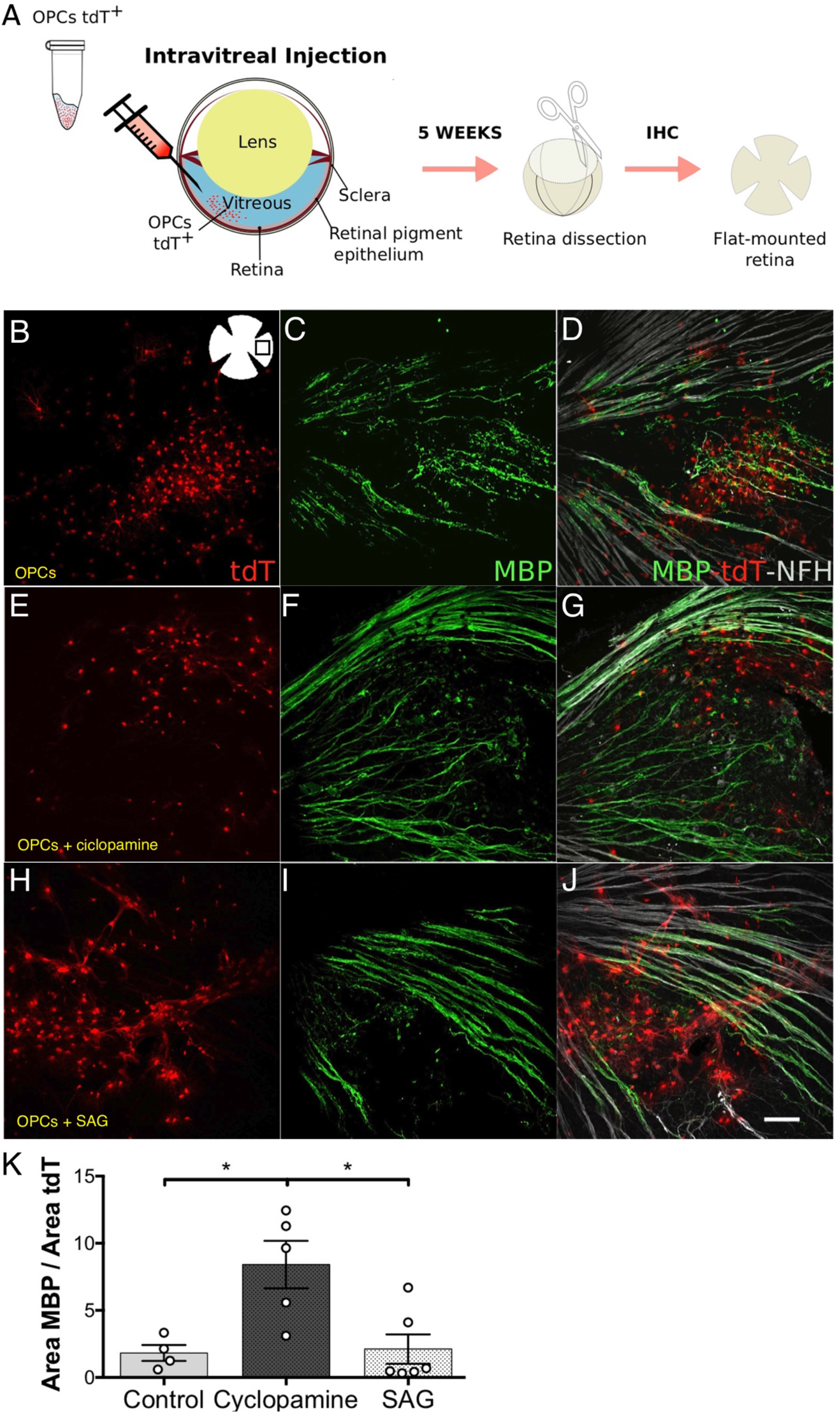
OPCs transplanted into the vitreous humor survive and retain their ability to differentiate into elements capable of myelination of retinal ganglion cell axons. Schematic of transvitreal retinal injection into the mouse eye and dissection for IHC (**A**). tdT^+^ cells (red) differentiate and form myelin sheaths (MBP, green) around axons of retinal ganglion cells (NFH, white) **(B-D)**. Coinjection of OPCs with cyclopamine results in increased amounts of myelin (**E-G**). Coinjection of OPCs with SAG does not lead to changes over what was observed in control animals (**H-J**). The quantification of the MBP area over the tdT^+^ element area reveals a significant increase in myelination in the group of animals treated with cyclopamine. Each circle on the graph corresponds to a retina. Statistical analysis was carried out using a one-way ANOVA test. **(K)**. *Scale bar: 100 μm*.

Additionally, to rigorously test our pharmacological manipulations of activating and inhibiting the Smo receptor, we injected cortical OPCs from the Smo^fl/fl^/NG2-Cre^ERT2^ (deletion of the Smo receptor), SmoM2/NG2-Cre^ERT2^ (activation of the Smo receptor) and NG2-Cre^ERT2^ (control) mice. Mice injected with control-OPCs (control) showed moderate myelination in retinal axons (supplementary figure 1). The myelinated area in the retina was markedly increased when OPCs from Smo^fl/fl^/NG2-Cre^ERT2^ mice were injected (supplementary figure 1). Finally, intravitreal injection of OPCs from the SmoM2/NG2-Cre^ERT2^ mice resulted in a moderate increase in myelinated area compared to the value of the control group (supplementary figure 1).

### During postnatal development, the activation of the Smo receptor negatively affects OPC differentiation

Shh is a mitogen, however, it is unclear whether this factor expands early progenitor cells or only those that are committed to a specific lineage. To assess the role of Shh during postnatal myelination, we examined the phenotype generated by knocking out the Smo receptor in OPCs in Smo^fl/fl^/NG2-Cre^ERT2^ mice and expressing a constitutive active form of Smo in OPCs with the SmoM2/NG2-Cre^ERT2^ mice during postnatal myelination of the corpus callosum at 14 days of age. Transgenics mice were administrated with tamoxifen at P1-P2 by the lactating mothers.

The tdT reporter was also crossed into the transgenic mice to allow for the identification of recombined cells. PDGFRα-positive recombined cells are classified as OPCs. Likewise, CC1-positive recombined cells are identified as mature oligodendrocytes (OLs) (figure 2A). The analysis revealed that the SmoM2/NG2-Cre^ERT2^ animals present a considerably higher number of OPCs (tdT^+^-PDGR*α*^+^) than the Smo^fl/fl^ and control mice (figure 2A-B). Likewise, no significant differences were observed between the Smo^fl/fl^/NG2-Cre^ERT2^ group and the control group in respect to the number of OPCs (figure 2A-B). When considering the population of OLs (tdT^+^-CC1^+^), SmoM2/NG2-Cre^ERT2^ animals displayed significantly lower numbers compared to both control and Smo^fl/fl^/NG2-Cre^ERT2^ groups (figure 2A-C). No significant differences were observed in the tdT^+^-CC1^+^ elements between control and Smo^fl/fl^ mice (figure 2A-C).

**Figure 2.**
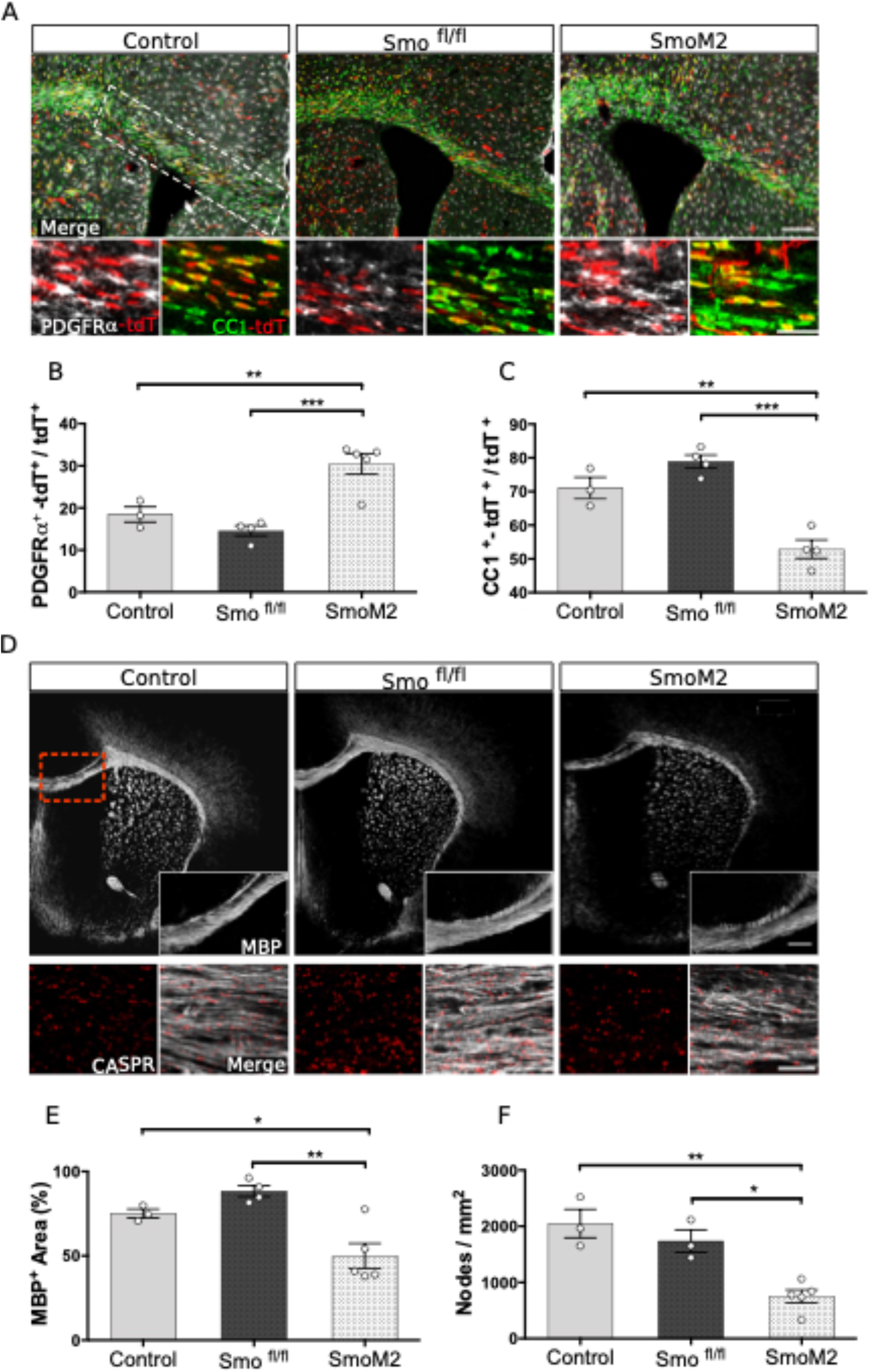
– Smo gain of function increases the rate of OPCs impeding their differentiation into mature OLs. Panoramic and detailed views of the CC in control, Smo^fl/fl^ and SmoM2 P14 mice injected with tamoxifen at P1-P2. White outlines highlight the region used for the quantification. Immunostaining for PDGFRα OPCs (white), CC1 mature OLs (green), and tdTomato (red). *Scale bar: 300 μm (100 μm in magnification)* (**A**). Quantification of the PDGFRα or CC1-tdT^+^ cells shows a higher rate of OPCs and lower OLs in SmoM2 mice. The statistical analysis was carried out using the one-way ANOVA test and the post-hoc Tukey’s multiple comparisons test, where **p< 0.01; ***p< 0.001. Each data point is an animal (**B-C**). Overview and detailed myelination of the rostral CC. Myelin stained with MBP (white) and paranodes with CASPR (red) showing that the increase of Smo receptor expression leads to a deficit in CNS myelination with a lower density of doublets. *Scale bar: 200 μm (30 μm in magnification)* (**D**). Quantification of MBP area and the number of CASPR doublets counted per mm^2^ show a decrease of myelin and paranode in the SmoM2 line. The statistical analysis was carried out using the one-way ANOVA test and the post-hoc Tukey’s multiple comparisons test, where *p< 0.05; **p< 0.01. Each data point is an animal (**E-F**).

In order to further analyze the impact that delayed differentiation of OLs has on myelination, we measured the area of myelin basic protein (MBP) and the number of nodes of Ranvier. The latter were calculated based on the number of CASPR doublets (axonal protein expressed at the paranode) per square millimeter. SmoM2 mice have significantly less MBP area in the corpus callosum than their Smo^fl/fl^ and control counterparts (figure 2D-E). In SmoM2 mice, the reduction in the presence of MBP reaches 24.71% in relation to the value of the control group; which is consistent with the 1.34-fold reduction in the number of OLs in the aforementioned SmoM2 mouse. In line with the above, the SmoM2 mice have a lower number of Ranvier’s nodes compared to the Smo^fl/fl^ and control lines (figure 2D, F). No differences were found in the organization of this axonal tracts between the groups (Supplemmentary Figure 2).

In view of these results, we can conclude that during development, the Shh/Smo pathway can regulate the differentiation of OPCs in the corpus callosum. Thus, Shh can positively influence the increase in the number of OPCs. However, Shh is not a necessary factor for myelination of the corpus callosum during postnatal development. If Shh/Smo is required during postnatal development, Smo^fl/fl^ mice would show marked differences from the control line, which is not observed.

### Constitutive activation of the Shh/Smo pathway promotes the proliferation of OPCs during active demyelination

Cuprizone induces the death of oligodendrocytes and, with it, demyelination after a few days of ingestion. Within the first three weeks of sustained cuprizone intake, demyelination, microglia activation, and OPC proliferation can be observed (figure 3A-I, 3M). This proliferation of OPCs begins at week 1-2 of treatment and reaches a peak after 3-4 weeks of cuprizone intake (Gudi *et al*., 2014). This part of the study was carried out on the corpus callosum after three weeks of cuprizone intake. In this experimental paradigm, animals were injected with tamoxifen for five days followed at the start of the cuprizone diet (figure 3A). To detect proliferating cells, EdU was dissolved in the drinking water after three weeks of cuprizone diet (figure 3A). This ensures that tdT^+^/EdU^+^ cells have proliferated during treatment with cuprizone.

**Figure 3.**
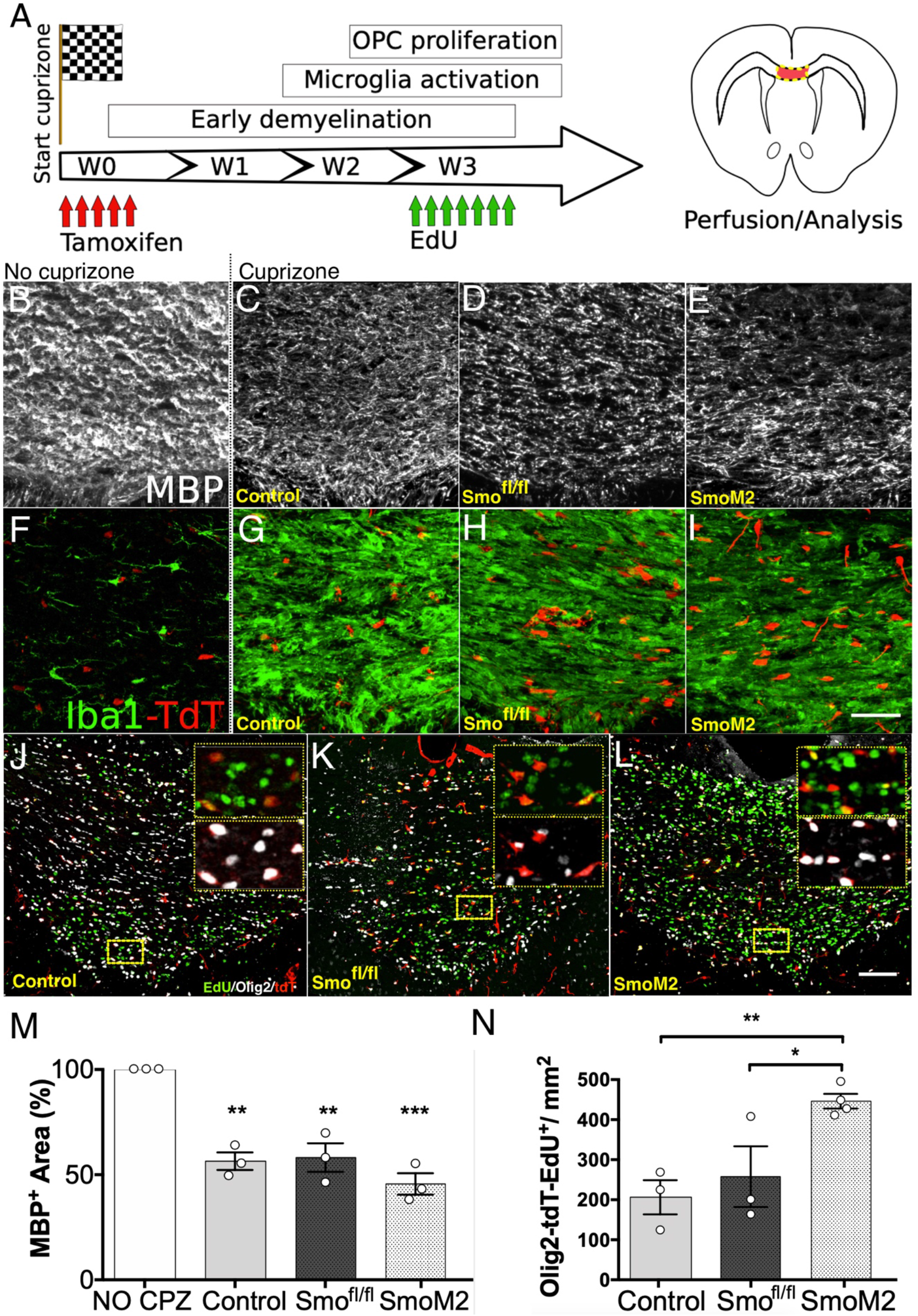
– In a context of active demyelination, overactivation of the Shh/Smo pathway leads to an increase in the proliferation of OPCs. Timeline scheme of the cuprizone-induced demyelination. Tamoxifen was injected with the beginning cuprizone diet, and EdU was administrated for seven consecutive days before the analysis (**A**). Dietary ingestion of cuprizone results in loss of myelin in treated animals (**B-E**). The presence of microglia (Iba1, green) is markedly higher in the region studied in NG2-Cre^ERT2^ (control), Smo^fl/fl^/NG2-Cre^ERT2^ and SmoM2/NG2-Cre^ERT2^ treated mice compared to that observed in the corpus callosum of animals not treated with cuprizone. *Scale bar: 50 mm* (**F-I**). Triple labeling with EdU (green), Olig2 (white) and the tdT reporter (red) in the corpus callosum of NG2-Cre^ERT2^ (control), Smo^fl/fl^/NG2-Cre^ERT2^ and SmoM2/NG2-Cre^ERT2^ animals. The boxes are the magnification of the marked area (yellow box) in the main image. *Scale bar: 100 mm* (**J-L**). The percentage of the area covered by myelin is significantly lower in all groups treated with cuprizone compared to the value obtained in untreated animals (**M**). Quantification of the density of triple labeled cells shows that the Shh pathway activation contributes to the increase of OPC proliferation in response to demyelination (**N**). The statistical analysis was carried out using the one-way ANOVA test and the post-hoc Tukey’s multiple comparisons test, where *p< 0.05, **p< 0.01 and ***p<0.001. Each data point is an animal.

During active demyelination, overactivation of the Smo pathway (SmoM2/NG2-Cre^ERT2^) leads to a significant increase in the proliferation of oligodendrocyte lineage cells (Olig2^+^/tdT^+^/EdU^+^ cells) compared to that observed in the control and Smo^fl/fl^ lines (figure 3J-L, N). Likewise, the loss of function (Smo^fl/fl^/NG2-Cre^ERT2^) did not result in significant changes with respect to control animals (figure 3J-L, N).

### Constitutive activation of the Shh/Smo pathway has no significant effect on oligodendroglial cells during remyelination

We next set out to evaluate the number of oligodendrocytes and myelination in the corpus callosum two weeks after the completion of cuprizone treatment, considered the remyelination phase (figure 4A). In these experiments, tamoxifen was administered to animals during the last week of cuprizone treatment (week 5) and the generation of the olig2^+^/tdT^+^ and CC1^+^/tdT^+^ cells were studied during the recovery phase (figure 4A).

**Figure 4.**
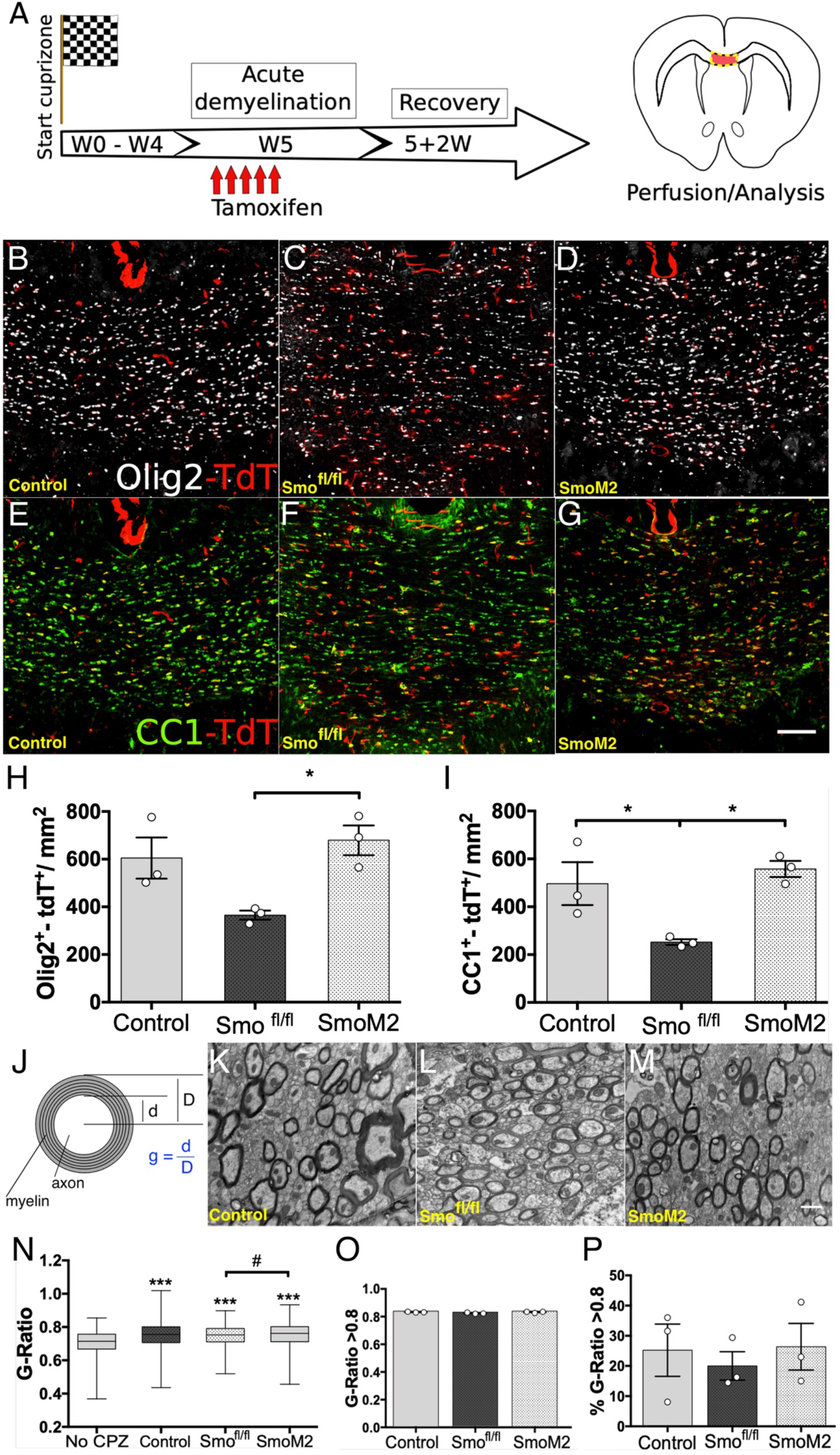
In a context of remyelination after injury, the loss of the Shh/Smo pathway results in a significant decrease in the number of OPCs and oligodendrocytes. Timeline scheme of the cuprizone-induced demyelination. Tamoxifen was injected during the last week of the cuprizone diet. Brains were analyzed after two weeks of recovery (**A**). Labeling of positive elements to Olig2 (white) and tdT (red) in the corpus callosum of control, Smo^fl/fl^/NG2-Cre^ERT2^ and SmoM2/NG2-Cre^ERT2^ mice (**B-D**). Labeling of positive elements to CC1 (green) and tdT (red) in control, Smo^fl/fl^/NG2-Cre^ERT2^ and SmoM2/NG2-Cre^ERT2^ mice (**E-G**). Quantification of the positive elements to Olig2 and CC1 in the experimental groups considered. In both cases, Smo^fl/fl^/NG2-Cre^ERT2^ (loss of the Shh/Smo pathway) mice present significantly lower values compared to their NG2-Cre^ERT2^ (control) and SmoM2/NG2-Cre^ERT2^ (overactivation of the Shh/Smo pathway) counterparts (**H-I**). G-ratio calculation (**J**). Electron microscopy of the corpus callosum of control, Smo^fl/fl^/NG2-Cre^ERT2^ and SmoM2/NG2-Cre^ERT2^ mice (**K-M**). Box and whisker plot shows an increase of the G-ratio of myelinated axons in cuprizone-treated mice with respect to the no-cuprizone group. The G-ratio in Smo^fl/fl^/NG2-Cre^ERT2^ mice presents a slightly lower mean value compared to its SmoM2/NG2-Cre^ERT2^ counterpart; although the differences do not reach statistical significance (**N**). There are no differences in the values of the G-ratio >0.8 between the three experimental groups (**O**). Quantification of the percentage of axons with a G-ratio greater than 0.8. The Smo^fl/fl^/NG2-Cre^ERT2^ mice present a slightly lower value in this percentage compared to the NG2-Cre^ERT2^ (control) and SmoM2/NG2-Cre^ERT2^ (overactivation of the Shh/Smo pathway) groups; although the differences are not statistically significant. There are no differences between the Smo2/NG2-Cre^ERT2^ and NG2-Cre^ERT2^ (control) groups (**P**). The statistical analysis was carried out using the one-way ANOVA test and the post-hoc Tukey’s multiple comparisons test, where *p< 0.05; **p< 0.01; ***p< 0.001. Each data point is an animal. *Scale bar: 100 mm (B-G) y 1 mm (K-M)*.

During remyelination, constitutive activation of the Shh/Smo pathway (SmoM2/NG2-Cre^ERT2^) did not translate into an increase in the Olig2^+^/tdT^+^ population compared to the control group (figure 4B,D,H). Likewise, no significant changes were observed in the number of mature oligodendrocytes (CC1^+^/tdT^+^) between the SmoM2 mice and the control line (Figure 4E,G,I). In contrast, loss of function (Smo^fl/fl^/NG2-Cre^ERT2^ mice) altered the number of cells of the oligodendroglial lineage. The Smo^fl/fl^ mice displayed a significantly lower number of CC1^+^/tdT^+^ cells compared to the control group (Figure 4E,F,I). Furthermore, although statistical significance was not achieved, there is an evident reduction of Olig2^+^/tdT^+^ cells in the Smo^fl/fl^/NG2-Cre^ERT2^ mice compared to the control group (NG2/Cre^ERT2^) (Figure 4B-C,H). In summary, the constitutive activation of the Shh/Smo pathway does not result in a significant change; but, loss of function negatively affects the number of cells of the oligodendrocyte lineage.

After quantification of the Olig2^+^/tdT^+^ and CC1^+^/tdT^+^ populations, we assessed the remyelination process by means of electron microscopy and the calculation of the g-ratio (figure 4J-P). A lower value in the g-ratio indicates a thicker myelin sheath (figure 4J; Murcia-Belmonte *et al*., 2016, Medina-Rodríguez *et al*., 2017). Analysis of the g-ratio reveals significant differences between animals treated and not treated with cuprizone (figure 4N). This validates the cuprizone treatment as a demyelinating agent prior to the remyelination phase. However, no differences were observed between the three experimental groups (control, SmoM2 and Smo^fl/fl^) (figure 4K-N). The g-ratio also allows discerning between remyelinated axons (g-ratio ≥0.8) and pre-existing myelinated axons (g-ratio < 0.8). We also did not find differences in this relationship between the three experimental groups (Figure 4O-P).

Table 1 summarizes the data that we obtained in the different experimental paradigms and the different myelinating or remyelinating scenarios.

**Table 1.** Changes observed in the four experimental paradigms considered. Ct: control conditions/groups; + : survival + differentiation; N: normal; High/Low: number of cells (related to Ct).

| *In vivo* paradigm | Type of cells | Smo<sup>fl/fl</sup>/NG2-Cre<sup>ERT2</sup><br>(low Shh) | Control<br>(Ct) | SmoM2/NG2-Cre<sup>ERT2</sup><br>(high Shh) |
| --- | --- | --- | --- | --- |
| **OPCs Intravitreal transplant** (adult)<br>Retinal Ganglion Cell Layer | **OPCs** | + | + | + |
|  | **OLs / Myelin** | +++ | + | + |
| **Development** (p14)<br>Corpus callosum | **OPCs** | N | N | High |
|  | **OLs / Myelin** | N | N | Low |
| **Demyelination** (adult)<br>Cuprizone | **OPCs** | N | N | High |
|  | **OLs / Myelin** | N | N | N |
| **Remyelination** (adult)<br>After cuprizone damage | **OPCs** | Low | N | N |
|  | **OLs / Myelin** | Low | N | N |

## DISCUSSION

The mechanisms underlying how Smo-mediated Shh signaling controls oligodendroglial differentiation in (re)myelination remain unclear due to the different experimental paradigms that result in inconsistent results, including the use of immortalized cell lines (Ruat *et al*., 2014; Del Giovane and Ragnini-Wilson, 2018, Dal Giovine *et al*., 2022). Our present work aims to clarify these differences by studying Smo-mediated Shh signaling in *i)* a naïve scenario for myelination, *ii)* a developmental scenario for myelination, iii) cuprizone-induced demyelination, and *iv)* remyelination after demyelination with cuprizone.

The retina offers an unbeatable framework for the study of the myelination process: a set of axons ready to be myelinated without interference. This experimental framework offers unquestionable advantages over other in vitro methods used to assess the effect of factors and molecules. Thus, the retina offers a three-dimensional structure and in vivo interactions that are difficult to recreate in vitro. However, we first checked if OPCs and OLs themselves can survive in a structure that OPCs do not colonize under normal conditions in most vertebrates (this particularity is reviewed in: Sugimoto *et al*., 2001; Spassky *et al*., 2002). Our present results confirmed that OPCs injected into the vitreous humor of young adult mice survive and retain their ability to proliferate and differentiate into myelin-forming phenotypes, even in the absence of concomitant exogenously induced inflammation (Setzu *et al*., 2006; Yuen *et al*., 2013). With this, we significantly improve the framework to evaluate the role of different molecules and/or receptors in the myelination process. Once the viability of OPCs within the vitreous was confirmed, we tested the inhibition of Smo receptor in the intravitreal transplanted OPCs, either pharmacologically (with cyclopamine) or genetically (transplanting OPCs isolated from the Smo^fl/fl^/NG2-Cre^ERT2^ mice). In both conditions, there is an evident increase in the amount of myelin that surrounds the axons of the retinal ganglion cells in relation to control mice. On the contrary, the stimulation of Smo pathway (either pharmacologically with SAG, or transplanting OPCs isolated from the SmoM2/NG2-Cre^ERT2^ transgenic mice) did not have effects. We therefore conclude that in a *naïve* environment (the adult retina), the inhibition of the Smo pathway leads to the differentiation of OPCs into mature OLs and thus significantly more myelination of the intraretinal segment of the retinal ganglion cell axons (Table 1).

But, as has just been stated, the adult retina is a “*naïve*” environment for myelin formation. It is free of developmental or reparative programs. What would happen in other environmental conditions? During postnatal development (P14), upregulation of the Shh/Smo pathway results in a significant increase in the number of OPCs, as well as a decrease in mature oligodendrocytes. In addition, the gain of function implied a decrease in the area of myelinated axons in the corpus callosum and a markedly decrease in the number of nodes of Ranvier (Table 1). Previous studies already pointed in this direction. Thus, it had been shown that *in vitro* (radial glia cell cultures) treatment with Shh induced an increase in the number of Olig2^+^ progenitors (Ortega *et al*., 2013). In addition, the spinal cord of SmoM2 mice displayed a higher number of OPCs without affecting their proliferation, together with lower degrees of myelination (Xu *et al*., 2020). In agreement with these authors, we suggest that Shh acting via Smo receptor is not required for terminal differentiation of OPCs, and these cells are present but “arrested” in this undifferentiated state in the SmoM2/NG2-Cre^ERT2^ mice (Table 1).

The presence of Shh has also been described in processes other than CNS development, like demyelination, including multiple sclerosis in humans (Wang *et al*., 2008; de Castro *et al*., 2013; Macchi *et al*., 2020). To further explore Shh-Smo signaling, we used cuprizone demyelination to study the role of overactivation and loss of the Shh-Smo pathway during an active demyelination process and the subsequent the remyelinating scenario after the damage process. In the active demyelination process, upregulation of the Shh/Smo pathway (SmoM2/NG2-Cre^ERT2^ mice) increases the number of Olig2^+^ cells compared to the control group (Table 1). Again, as in the development of the CNS, overactivation of the Shh-Smo pathway is sufficient to maintain OPCs in an undifferentiated state.

The remyelination phase after cuprizone damage is revealed as a new scenario that does not seem to be the simple continuation of what happens during the active period of demyelination. In the remyelination scenario, upregulation of the Shh/Smo pathway (SmoM2/NG2-Cre^ERT2^) does not translate into an increase in cells of the oligodendrocyte lineage (Olig2^+^ and CC1^+^) compared to control (NG2-Cre^ERT2^). During the remyelination phase, the factors that have potentiated the mitogenic character of Shh in the active demyelination may no longer be present. On the other hand, the loss of the Shh/Smo pathway (Smo^fl/fl^/NG2-Cre^ERT2^) during the remyelination phase implies a reduction in the number of mature oligodendrocytes compared to the control and SmoM2 groups. With this, it would be expected that myelination observed in the Smo^fl/fl^ animals would also be attenuated in comparison to the control and SmoM2 groups. That is, the less mature oligodendrocytes, the less myelin. However, this is not the case. Despite having fewer mature oligodendrocytes, Smo^fl/fl^ mice have the same g-ratio values and g-ratio > 0.8 percentage as control and SmoM2. It could be said that there are fewer mature oligodendrocytes, but these myelinate more. This is not surprising if we reconsider the data obtained on the myelination of retinal ganglion cell axons. In both the retina and the remyelination phase, mature oligodendrocytes independent of the Shh/Smo pathway myelinate more.

Altogether, our results suggest that the modulation of Shh signaling depends on the environment. During *i)* development, Smo mediated signaling cooperates with other growth factors to promote OPC proliferation (Kessaris *et al*., 2004; Furusho *et al*., 2011). Once OPCs reach a suitable density, the pathway has to be inhibited in favor of the differentiation. During the *ii)* demyelination phase, the environment is surrounded by proinflammatory factors. Here Shh acts mainly as a mitogen and could be helpful for the early stage OPCs to promote their proliferation and subsequently enhance the number of available remyelinating cells (Traiffort *et al*., 2020). However, Shh does not help these OPCs differentiate towards myelin-forming phenotypes (OLs), as proliferation and differentiation are mutually exclusive processes. When proinflammatory factors are downregulated during the *iii)* remyelination phase and OPCs have reached the appropriate density, inhibition of the Shh/Smo pathway may favor the remyelination process.

Ultimately, Shh/Smo upregulation is sufficient to maintain OPCs in an undifferentiated and proliferative state; but this same pathway is not necessary for the appearance of differentiated mature oligodendrocytes with the capacity to myelinate.

## Acknowledgments

*This research was supported by the Spanish Ministerio de Ciencia e Investigación-MCIN/Spanish Research* Agency-AEI grants SAF2012-400232, SAF2016-77575-R, RD12-0032/0012, RD16-0015/0019, *PID2019-109858RB-I00*, EIN2020-112366, *CSIC/Spanish Research Council grants BDC-*20213231 and 2023AEP096, and Fundación Ramón Areces (Spain) grant CIVP19A5917, and grant from Proyectos de Investigación Relacionados con las Enfermedades Raras Infantiles from Fundación Inocente Inocente (Spain), all to F.dC.

SN had a predoctoral contract of the Universidad de Castilla-La Mancha, and then was hired under grant IND2018/BMD-9751 (Comunidad de Madrid, Spain) to F.dC. BF-G was predoctoral researcher contracted under IND2018/BMD-9751 (Programa de Doctorados industriales1 de la Comunidad de Madrid, Spain), to F.dC. YL was hired under grants from Spanish Ministerio de Ciencia e Investigación-MCIN/Spanish Research Agency-AEI RD16/0015/0019 (ReTics Program, Instituto de Salud Carlos III) and *PID2019-109858RB-I00*.

## FIGURE LEGENDS

**Supplementary Figure 1.**
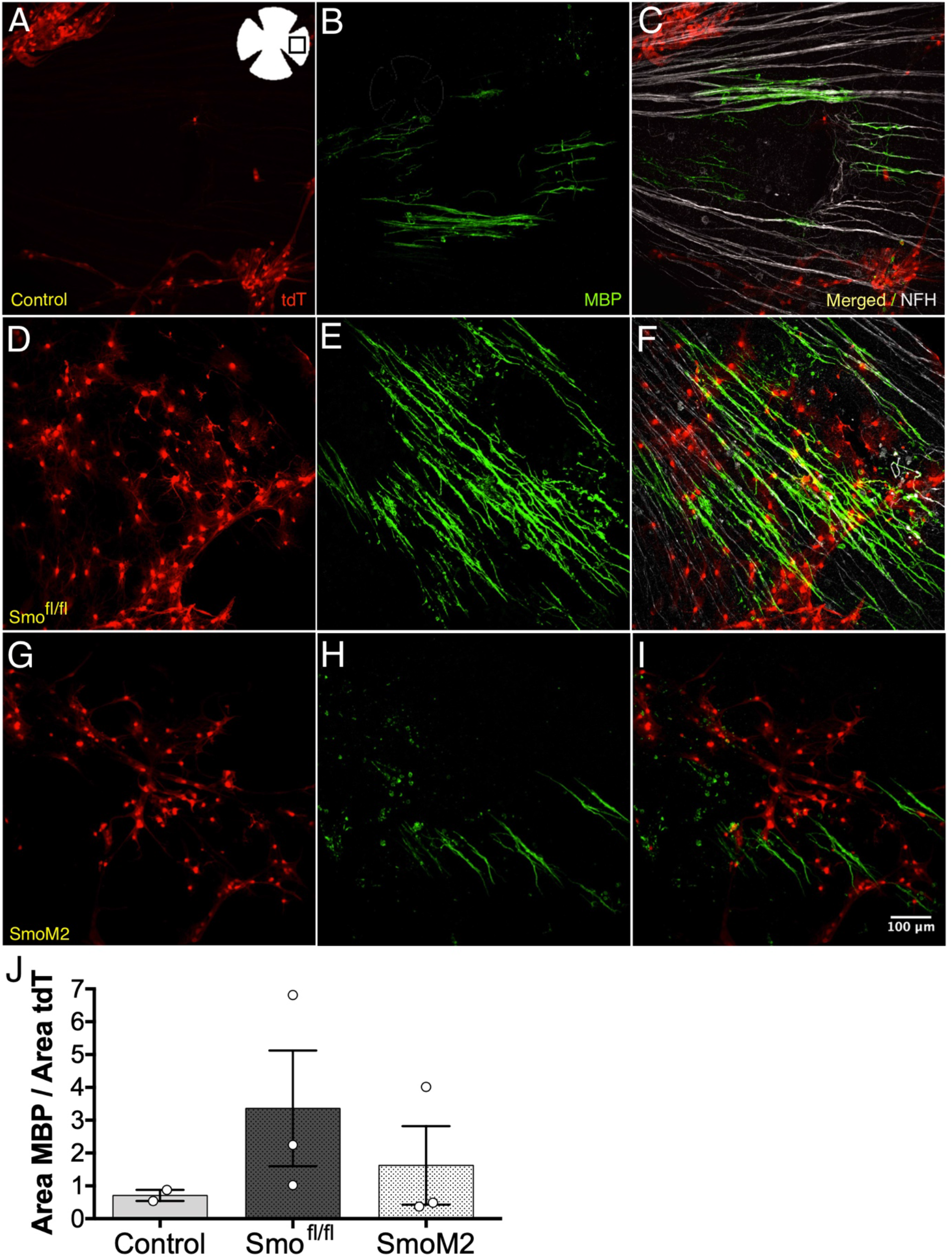
Intravitreal injection of tdT^+^ OPCs from three transgenic lines reveals greater myelination in the group without Smo function. Expression of tdT (red), MBP (green) and NFH (white) in retinas injected with OPCs from p7 NG2-Cre^ERT2^ animals (control group) (**A-C**). Expression of tdT (red), MBP (green), and NFH (white) in retinas injected with OPCs from p7 Smo^fl/fl^/NG2-Cre^ERT2^ animals (Smo loss-of-function) **(D-E)**. Expression of tdT (red), MBP (green), and NFH (white) in retinas injected with OPCs from p7 SmoM2/NG2-Cre^ERT2^ animals (Smo gain-of-function) (**G-I**). Quantification of the myelinated area (MBP area) in relation to the area with tdT^+^ elements present. The retinas of animals injected with OPCs from Smo^fl/fl^/NG2-Cre^ERT2^ animals show the highest value of myelinated area (**J**). *Scale bar: 100 μm*.

**Supplementary Figure 2.**
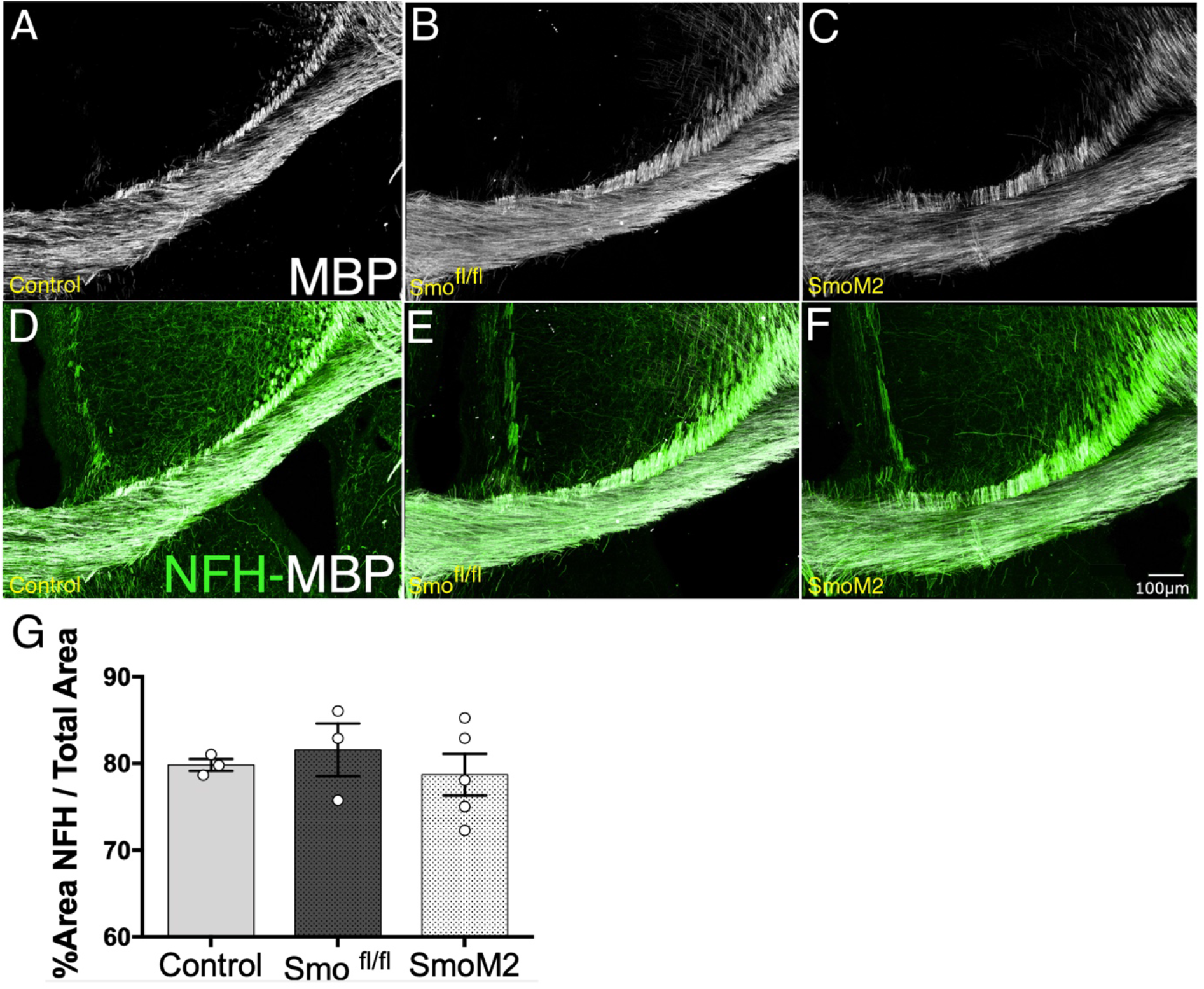
Distribution of NFH in the developing corpus callosum of the experimental groups considered. Myelin labeling (MBP, white) in the developing corpus callosum of control, Smo^fl/fl^, and SmoM2 mice (**A-C**). Double immunostaining of myelin (MBP, white) and axons (NFH, green) in the retinas of the three experimental groups considered (**D-F**). The NFH area with respect to the total area is quite similar in all the groups analyzed; with only a very slight increase in Smo^fl/fl^ mice compared to their control and SmoM2 counterpart (**G**).

## Notes

**Conflict of interest statement** The authors declare no conflicts of interest.

**Data availability statement** The data that support the findings of this study are available from the corresponding author upon reasonable request.

### Competing Interest Statement

The authors have declared no competing interest.

## REFERENCES

1. P. del Río-Hortega, El tercer elemento de los centros nerviosos. Poder fagocitario y movilidad de la microglía. Bol. de la Soc. Esp. de Biol. IX, (1919).

2. R.K. Small, P. Riddle, M. Noble, Evidence for migration of oligodendrocyte—type-2 astrocyte progenitor cells into the developing rat optic nerve. Nature 328(6126), 155–7 (1987).

3. G. Wolswijk, M. Noble, Identification of an adult-specific glial progenitor cell. Development 105(2), 387–400 (1989).

4. B.C. Warf, J. Fok-Seang, R.H. Miller, Evidence for the ventral origin of oligodendrocyte precursors in the rat spinal cord. J. Neurosci. 11(8), 2477–88 (1991).

5. J.M. Levine, R. Reynolds, J.W. Fawcett, The oligodendrocyte precursor cell in health and disease. Trends Neurosci. 24(1), 39–47 (2001).

6. R.J.M. Franklin, C. ffrench-Constant, J.M. Edgar, K.J. Smith, Neuroprotection and repair in multiple sclerosis. Nat. Rev. Neurol. 8(11), 624–34 (2012).

7. M. Fuccillo, A.L. Joyner, G. Fishell, Morphogen to mitogen: the multiple roles of hedgehog signalling in vertebrate neural development. Nat. Rev. Neurosci. 7(10), 772–83 (2006).

8. Y. Wang, J. Imitola, S. Rasmussen, K.C. O’Connor, S.J. Khoury, Paradoxical dysregulation of the neural stem cell pathway sonic hedgehog-Gli1 in autoimmune encephalomyelitis and multiple sclerosis. Ann. Neurol. 64(4), 417–27 (2008).

9. J. Ferent, C. Zimmer, P. Durbec, M. Ruat, E. Traiffort, Sonic Hedgehog signaling is a positive oligodendrocyte regulator during demyelination. J. Neurosci. 33(5), 1759–72 (2013).

10. A.J. Mierzwa G.M. Sullivan, L.A. Beer, S. Ahn, R.C. Armstrong, Comparison of cortical and white matter traumatic brain injury models reveals differential effects in the subventricular zone and divergent Sonic hedgehog signaling pathways in neuroblasts and oligodendrocyte progenitors. ASN Neuro. 6(5), p(2014).

11. J. Samanta et al., Inhibition of Gli1 mobilizes endogenous neural stem cells for remyelination. Nature 526(7573), 448–52 (2015).

12. A. Del Giovane, A. Ragnini-Wilson, Targeting Smoothened as a new frontier in the functional recovery of central nervous system demyelinating pathologies. Int. J. Mol. Sci. 19(11), 3677 (2018).

13. M.A. Sanchez, R.C. Armstrong, Postnatal Sonic hedgehog (Shh) responsive cells give rise to oligodendrocyte lineage cells during myelination and in adulthood contribute to remyelination. Exp. Neurol. 299, 122–136 (2018).

14. C.C. Winkler et al., The dorsal wave of neocortical oligodendrogenesis begins embryonically and requires multiple sources of Sonic hedgehog. J. Neurosci. 38(23), 5237–5250 (2018).

15. P. Namchaiw et al., Temporal and partial inhibition of Gli1 in neural stem cells (NCSs) results in the early maturation of NSC derived oligodendrocytes in vitro. Stem Cell Res. Ther. 10(1), 272 (2019).

16. M. Zakaria et al., The Shh receptor Boc is important for myelin formation and repair. Development. 146(9), dev172502 (2019).

17. M. Macchi et al., Mature oligodendrocytes bordering lesions limit demyelination and favor myelin repair via heparan sulfate production. Elife 9, e51735 (2020).

18. X. Ming, J.L. Dupree, V. Gallo, L.J. Chew, Sox17 promotes oligodendrocyte regeneration by dual modulation of hedgehog and Wnt signaling. iScience. 23(10), 101592 (2020).

19. A. Del Giovane et al., Smoothened/AMP-activated protein kinase signaling in oligodendroglial cell maturation. Front Cell Neurosci. 15, 801704 (2021).

20. J.A. Klein et al., Sonic hedgehog pathway modulation normalizes expression of Olig2 in rostrally patterned NPCs with trisomy 21. Front Cell Neurosci. 15, 794675 (2021).

21. Y. Laouarem et al., Functional cooperation of the hedgehog and androgen signaling pathways during developmental and repairing myelination. Glia. 69(6), 1369–1392 (2021).

22. V. Nguyen et al., Neuroprotective effects of Sonic hedgehog agonist SAG in a rat model of neonatal stroke. Pediatr. Res. 90(6), 1161–1170 (2021).

23. R. Petrova, A.L. Joyner, Roles for Hedgehog signaling in adult organ homeostasis and repair. Development. 141(18), 3445–57 (2014).

24. M. Ruat, L. Hoch, H. Faure, D. Rognan, Targeting of Smoothened for therapeutic gain. Trends Pharmacol Sci. 35(5), 237–46 (2014).

25. K. Loulier, M. Ruat, E. Traiffort, Increase of proliferating oligodendroglial progenitors in the adult mouse brain upon Sonic hedgehog delivery in the lateral ventricle. J. Neurochem. 98(2), 530–42 (2006).

26. Y. Xiao et al., Targeting central nervous system extracellular vesicles enhanced triiodothyronine remyelination effect on experimental autoimmune encephalomyelitis. Bioact. Mater. 9, 373–384 (2021).

27. D.Z. Radecki et al., Relative levels of Gli1 and Gli2 determine the response of ventral neural stem cells to demyelination. Stem Cell Reports. 15(5), 1047–1055 (2020).

28. N. Kessaris, F. Jamen, L.L. Rubin, W.D. Richardson, Cooperation between sonic hedgehog and fibroblast growth factor/MAPK signalling pathways in neocortical precursors. Development. 131(6), 1289–98 (2004).

29. F. de Castro, A. Bribián, The molecular orchestra of the migration of oligodendrocyte precursors during development. Brain Res. Brain Res. Rev. 49(2), 227–41 (2005).

30. M. Furusho, Y. Kaga, A. Ishii, J.M. Hébert, R. Bansal, Fibroblast growth factor signaling is required for the generation of oligodendrocyte progenitors from the embryonic forebrain. J. Neurosci. 31(13), 5055–5066 (2011).

31. F. de Castro, B. Zalc, Migration of myelin-forming cells in the CNS. Cellular Migration and Formation of Axons and Dendrites, 515–529 (2020).

32. V. Gudi, S. Gingele, T. Skripuletz, M. Stangel, Glial response during cuprizone-induced de- and remyelination in the CNS: Lessons learned. Front Cell Neurosci. 8:73 (2014).

33. V. Murcia-Belmonte et al., Anosmin-1 over-expression regulates oligodendrocyte precursor cell proliferation, migration and myelin sheath thickness. Brain Struct Funct. 221(3), 1365–1385 (2016).

34. E.M. Medina-Rodríguez et al., Promoting in vivo remyelination with small molecules: A neuroreparative pharmacological treatment for Multiple Sclerosis. Sci Rep. 7:p 43545 (2017).

35. Y. Sugimoto et al., Guidance of glial precursor cell migration by secreted cues in the developing optic nerve. Development. 128(17), 3321–30 (2001).

36. N. Spassky et al., Directional Guidance of Oligodendroglial Migration by Class 3 Semaphorins and Netrin-1. J Neurosci. 22(14), 5992–6004 (2002).

37. A. Setzu et al., Inflammation stimulates myelination by transplanted oligodendrocyte precursor cells. Glia. 54(4), 297–303 (2006).

38. T.J. Yuen et al., Identification of endothelin 2 as an inflammatory factor that promotes central nervous system remyelination. Brain. 136(4), 1035–47 (2013).

39. J.A. Ortega, N.V. Radonjić, N. Zecevic, Sonic hedgehog promotes generation and maintenance of human forebrain Olig2 progenitors. Front Cell Neurosci. 7:254 (2013).

40. X. Xu et al., Stage-specific regulation of oligodendrocyte development by Hedgehog signaling in the spinal cord. Glia. 68(2), 422–434 (2020).

41. F. de Castro, A. Bribián, M.C. Ortega, Regulation of oligodendrocyte precursor migration during development, in adulthood and in pathology. Cell Mol Life Sci. 70(22), 4355–4368 (2013).

42. T. Gorojankina et al., Discovery, molecular and pharmacological characterization of GSA-10, a novel small-molecule positive modulator of Smoothened. Mol. Pharmacol. 83, 1020–1029 (2013).

43. A. Fleury et al., Hedgehog associated to microparticles inhibits adipocyte differentiation via a non-canonical pathway. Sci Rep. 6:23479 (2016).

44. T. Akhshi, W.S. Trimble, A non-canonical hedgehog pathway initiates ciliogenesis and autophagy. J. Cell Biol. 220(1):e202004179 (2021).

45. J. Zhang et al., Fingolimod treatment promotes proliferation and differentiation of oligodendrocyte progenitor cells in mice with experimental autoimmune encephalomyelitis. Neurobiol Dis. 76:57–66 (2015).

46. Y. Wang, J.F. Martin, C.B. Bai, Direct and indirect requirements of Shh/Gli signaling in early pituitary development. Dev Biol. 348(2), 199–209 (2010).

47. G. Porcu et al., Clobetasol and halcinonide act as smoothened agonists to promote myelin gene expression and RxRγ receptor activation. PLoS ONE. 10(12):e0144550 (2015).

48. F.J. Najm et al., Drug-based modulation of endogenous stem cells promotes functional remyelination in vivo. Nature. 522(7555), 216–220. (2015).

49. E. Nocita et al., EGFR/ERBB inhibition promotes OPC maturation up to axon engagement by co-regulating PIP2 and MBP. Cells. 8(8), 844 (2019).

50. F. Long, X.M. Zhang, S. Karp, Y. Yang, A.P. McMaho, Genetic manipulation of hedgehog signaling in the endochondral skeleton reveals a direct role in the regulation of chondrocyte proliferation. Development. 128(24), 5099–108 (2001).

51. J. Jeong, J. Mao, T. Tenzen, A.H. Kottmann, A.P. McMahon, Hedgehog signaling in the neural crest cells regulates the patterning and growth of facial primordia. Genes Dev. 18(8), 937–951 (2004).

52. N. Kessaris, M. Fogarty, P. Iannarelli, M. Grist, M. Wegner, W.D. Richardson, Competing waves of oligodendrocytes in the forebrain and postnatal elimination of an embryonic lineage. Nat Neurosci. 9(2), 173–179 (2006).

53. T.A. Dincman, J.E. Beare, S.S. Ohri, S.R. Whittemore, Isolation of cortical mouse oligodendrocyte precursor cells. J. Neurosci. Methods, 209(1), 219–226 (2012).

54. A. Bribián et al., Functional heterogeneity of mouse and human brain OPCs: Relevance for preclinical studies in multiple sclerosis. J. Clin. Med. 9(6), 1–21 (2020).

55. J. Schindelin et al., Fiji: an open-source platform for biological-image analysis. Nat Methods. 9(7), 676–82 (2012).

